# Metabolic Versatility of *Lysinibacillus capsici* BCSIR-Raj-01 Underpins Its Multifunctionality in Biocontrol, Bioremediation, and Plant Growth Promotion

**DOI:** 10.1101/2025.10.09.681324

**Authors:** Robel H. Patwary, Md. Zamilur Rahman, Nahid Sultana, Natasha Nafisa Haque, Sanjana Fatema Chowdhury, Showti Raheel Naser, Mousona Islam, Tanay Chakrovarty, Md. Khalid Hasan, Ema Akter, Md. Abdullah Al Obaid, Md. Ahsan Habib, Shahina Akter, Tanjina Akhtar Banu, Barna Goswami, Md Murshed Hasan Sarker

## Abstract

Microbial inoculants that integrate plant growth promotion, pathogen suppression, and pollutant degradation are central to sustainable agriculture. Here, we report the genomic and phenotypic characterization of *Lysinibacillus capsici* BCSIR-Raj-01, an endophyte isolated from citrus leaves in Bangladesh. The 4.62 Mb genome encodes 4,699 coding sequences and displays an open pangenome with loci linked to stress adaptation, xenobiotic degradation, and secondary metabolite biosynthesis. Genome mining revealed complete pathways for indole-3-acetic acid (IAA) production, siderophore biosynthesis, and phosphate metabolism, alongside gene clusters for bacteriocins, lantibiotics, terpenes, and polyketides. Nitrogen cycling genes indicated diazotrophic growth potential and a truncated denitrification pathway. Phenotypic assays validated these predictions: *L. capsici* BCSIR-Raj-01 solubilized phosphate, produced IAA and siderophores, grew on hydrocarbons, and exhibited diazotrophic activity. Notably, the strain showed potent larvicidal activity against *Culex* and *Anopheles* larvae, suggesting ecological roles beyond plant association. Together, these results establish *L. capsici* BCSIR-Raj-01 as a multifunctional bacterium with potential applications in crop improvement, bioremediation, and biological vector control.

**IMPORTANCE:** The challenges of modern agriculture and environmental sustainability demand microbial solutions that deliver multiple functions. *Lysinibacillus* species are increasingly recognized for such versatility, but their genomic basis remains poorly resolved. Our study provides the first genome-enabled framework for understanding multifunctionality in *L. capsici*. By combining whole-genome sequencing with phenotypic validation, we show that BCSIR-Raj-01 harbors biosynthetic clusters, stress adaptation loci, and xenobiotic degradation genes while also expressing experimentally confirmed traits, including phosphate solubilization, IAA and siderophore production, nitrogen fixation, hydrocarbon utilization, and larvicidal activity. This integration of plant growth promotion, biocontrol potential, and pollutant degradation positions *L. capsici* BCSIR-Raj-01 as a next-generation bioinoculant. Beyond broadening the functional scope of *Lysinibacillus*, our findings highlight how genome-guided approaches can identify microbes with cross-sectoral applications in agriculture, bioremediation, and vector management.

## INTRODUCTION

Modern agriculture and environmental management face a dual challenge: the intensification of chemical inputs to sustain yields, and the ecological costs of those inputs, from pesticide resistance to soil degradation and greenhouse gas emissions. There is a growing demand for microbial inoculants that can simultaneously promote plant growth, suppress pathogens, and mitigate environmental contamination (1). This multi-pronged challenge has positioned multifunctional bioinoculants—specifically those capable of concurrent plant growth promotion, pathogen suppression, and pollutant biodegradation—as a central goal in sustainable agriculture (2–4).

Within this context, the genus *Lysinibacillus* has emerged as a promising yet underexplored group of Gram-positive, endospore-forming bacteria. Originally reclassified from *Bacillus* based on cell wall peptidoglycan structure (5), *Lysinibacillus* species have been found in agricultural soils, contaminated sites, and the rhizosphere (6). Their ecological breadth hints at metabolic versatility. Indeed, *L. sphaericus* is well known for its entomopathogenic properties through Bin and Mtx toxins active against mosquito larvae (7), while other strains display classic plant growth-promoting (PGP) traits such as phosphate solubilization, siderophore production, indole-3-acetic acid (IAA) biosynthesis, and stress tolerance (8, 9).

Recent work extends the functional scope of *Lysinibacillus* beyond crop productivity. Certain isolates antagonize fungal phytopathogens via no n-ribosomal peptide and lipopeptide biosynthetic clusters (9, 10), while others degrade polycyclic aromatic hydrocarbons and synthetic polymers, underscoring their value in bioremediation (12). *Lysinibacillus capsici* in particular has drawn attention: strain TT41 was shown by metabolomic profiling to enhance drought tolerance in *Brassica rapa* through reprogramming host metabolism (13), and other isolates have inhibited soil-borne pathogens including *Rhizoctonia solani* and *Fusarium oxysporum*. Such multifunctionality positions *L. capsici* as a candidate at the intersection of agriculture and environmental sustainability.

Despite these advances, the genomic underpinnings of multifunctionality in *Lysinibacillus* remain poorly resolved. Genome-scale metabolic models (GEMs) have proven powerful for linking genotype to phenotype in related *Bacillus* and *Paenibacillus* strains, revealing determinants of rhizosphere competence and biocontrol activity (14, 15). Comparable genome-enabled frameworks are largely missing for *Lysinibacillus*. This knowledge gap limits our ability to harness its full potential as a bioinoculant.

Here, we address this gap by presenting the integrated genomic and phenotypic characterization of *Lysinibacillus capsici* BCSIR-Raj-01, an endophyte isolated from citrus leaves in Bangladesh. Through whole-genome sequencing, comparative genomics, and functional assays, we uncover the strain’s metabolic repertoire, stress tolerance mechanisms, and secondary metabolite biosynthetic potential. Experimental validation of plant growth–promoting traits, hydrocarbon degradation, and larvicidal activity complements the genomic predictions. Together, these findings provide a blueprint for positioning *L. capsici* BCSIR-Raj-01 as a multifunctional bacterium with applications in sustainable crop production, bioremediation, and biological vector control.

## MATERIALS AND METHODS

### Sample Collection and Isolation of *Lysinibacillus capsici* BCSIR-Raj-01

An endophytic bacterial strain, *Lysinibacillus capsici* BCSIR-Raj-01, was isolated from healthy citrus (*Citrus limon*) leaf tissues collected in Rajshahi, Bangladesh. Leaves were surface-sterilized using 70% ethanol and 1% sodium hypochlorite, followed by rinsing in sterile distilled water (16). The plant tissue is then cut into small pieces, and plated onto nutrient agar plates. After at 30°C for 48 h incubation, visible bacterial colonies are subcultured onto fresh media to obtain pure cultures for further characterization and analysis. Cultures were stored at −80°C in 25% glycerol.

### Genomic DNA extraction and whole-genome sequencing

Genomic DNA was extracted using the Qiagen Genomic DNA Kit (Qiagen, USA) and quantified by NanoDrop spectrophotometry (Thermo Scientific™ NanoDrop™ One Microvolume UV Spectrophotometer, USA). DNA libraries were prepared using the Nextera DNA Flex Library Preparation Kit (Illumina, San Diego, CA) following the manufacturer’s instructions. The resulting DNA libraries were purified using AMPure XP beads (Beckman Coulter, Sharon Hill, PA) and re-quantified using the Quantus™ Fluorometer (Promega). Sequencing was performed on the MiniSeq System using v2 sequencing reagent kits (Illumina). Read quality was checked using FastQC v0.11.9 and trimming was done using Trimmomatic v0.39 (17).

### Genome Assembly and Annotation

Genome assembly was carried out using SPAdes v3.15.3 (18) within the EDGE Bioinformatics platform (19). Assembly metrics were assessed with QUAST (20). Functional annotation was performed using the BV-BRC pipeline (21), supplemented with RAST (22), GO, KEGG (23), and EC number predictions. Genomic features were visualized using the Proksee platform (24).

### Pan-Genome and Phylogenomic Analysis

A total of 25 publicly available *Lysinibacillus* genomes were retrieved from NCBI GenBank. Pan-and core-genome analyses were conducted using the Integrated Prokaryotic Genome and Pan-genome Analysis (IPGA v1.09) platform (25). Phylogenetic trees were constructed with Type (Strain) Genome Server ( Meier-Kolthoff et al., 2019) using Genome BLAST Distance Phylogeny (GBDP) method.

### Secondary metabolite biosynthesis gene clusters

Secondary metabolite biosynthetic gene clusters (BGCs) were predicted using antiSMASH v5.0 (27), BAGEL4 (28), and the PATRIC secondary metabolite module (21).

### Mobile genetic elements and defense mechanisms

MobileOG-db (29) was used to detect mobile genetic elements (MGEs), classified by functional roles. Prophage regions were identified using PHAST (30). Restriction-modification systems were annotated with DefenseFinder (31, 32), CRISPR arrays were detected with CRISPRCasFinder (33), and other defense systems were analyzed via PADLOC (34).

### Xenobiotic biodegradation and resistance Genes

Xenobiotic degradation genes were identified using the HADEG database (35), including genes involved in aerobic hydrocarbon degradation. The BacMet database (36) was used to identify metal and biocide resistance genes. CARD (37) and ResFinder (38) were used for profiling antibiotic resistance genes. Genomic mapping and visualization were carried out using PROKSEE (24).

### Stress response and environmental adaptation

Stress-related genes were annotated using the PATRIC stress response module (21), and classified via KEGG and COG. Genes related to oxidative stress, osmotic stress, heat shock, and general stress adaptation were documented.

### Functional assays for plant gowth-promoting traits

Phosphate solubilization was evaluated on Pikovskaya’s agar medium, which contains insoluble tricalcium phosphate as the primary phosphate source (39). The isolate was spot-inoculated and incubated at 30 °C for 7 days. Solubilization activity was indicated by the formation of a clear halo surrounding bacterial colonies, and the solubilization index (SI) was calculated as the ratio of the total halo diameter to colony diameter (40, 41). Siderophore production was determined using chrome azurol S (CAS) agar assay, which detects the removal of Fe³⁺ from the CAS-dye complex (42). A color change from blue to orange around colonies was considered positive for siderophore production. The intensity of color change was visually compared with uninoculated controls, and qualitative results were recorded.

Indole-3-acetic acid (IAA) production was quantified using the Salkowski colorimetric method (43, 44). Briefly, the isolate was cultured in Luria–Bertani (LB) broth supplemented with 0.5 g/L L-tryptophan at 30 °C for 48 h under shaking conditions (150 rpm). Cultures were centrifuged at 10,000 × g for 10 min, and 1 mL of cell-free supernatant was mixed with 2 mL of freshly prepared Salkowski reagent (1 mL of 0.5 M FeCl₃ in 50 mL of 35% HClO₄). IAA production was confirmed by the visual shift in color from yellow to red after 30 minutes of incubation in the dark. The ability of *L. capsici* BCSIR-Raj-01 to fix atmospheric nitrogen was assessed on nitrogen-free Jensen’s agar medium (45, 46). The strain was spot-inoculated onto plates containing sucrose as the sole carbon source and incubated at 30 °C for 5–7 days. Nitrogen fixation was indicated by visible bacterial growth in the absence of combined nitrogen sources. Acid production resulting from carbohydrate utilization was detected as a yellow zone surrounding colonies due to the pH indicator incorporated in the medium.

### Hydrocarbon degradation assay

The hydrocarbon-degrading potential of the *L. capsici* BCSIR-Raj-01 was established using Bushnell-Haas (BH) medium, which has inorganic salts but not an organic carbon source and thus ensured diesel to be the sole source of carbon. Diesel fuel was sterilized through filtration (0.22 μm) prior to use. The assays were carried out in aseptic 96-well plates in which 100 µL of 2× BH broth was put into each well, to which 25 µL or 50 µL of diesel and 50 µL of bacterial suspension (OD600 ≈ 0.1) was added. Each well volume was brought to 200 µL, and all the treatments were carried out in triplicate. Controls included (i) BH medium and diesel but no bacterial inoculation and (ii) diesel alone. Aerobic incubation was carried out at 30 °C for 7 days under static conditions. Bacterial growth was monitored spectrophotometrically (SpectraMax iD5 Multi-Mode Microplate Reader, USA) by determining the 600 nm (OD600) optical density at 24 h, 48 h, and daily up to date 7. An increase in OD600 compared to controls was taken as a parameter for diesel utilization by the isolate.

### Larvicidal activity against *Culex pipiens*

The larvicidal activity of *L. capsici* BCSIR-Raj-01 was evaluated against third-instar *Culex pipiens* larvae following WHO guidelines (47). Bacterial cultures were grown at 30 °C to ∼10⁹ CFU/mL, and aliquots (10–200 µL) were added to 100 mL bioassay solutions. Each concentration was tested in triplicate, with 20 larvae maintained in 30 mL of test solution per sterile beaker . All the assays were conducted under controlled environmental conditions (30 °C, 70–75% RH, 12:12 light–dark cycle) without larval feeding. The mortality was checked at 24 h and 48h and larvae were considered dead if they did not respond to slight poking. Mortality values were corrected for using Abbott’s formula (48) and analyzed using Probit regression via SPSS v25 to derive LC₅₀ and LC₉₀ with 95% confidence intervals (49).

### Statistical Analysis

All experiments were conducted in triplicate. Results are expressed as mean ± standard deviation (SD). Statistical significance was assessed via one-way ANOVA followed by Tukey’s post hoc test using R Studio version 4.0.2 (R Core Team, 2022). A *p* value < 0.05 was considered significant.

## RESULTS

### Genome features and comparative genomics of *Lysinibacillus capsici* BCSIR-Raj-01

#### Genome Assembly and Phylogenetic analysis

The genome of *Lysinibacillus capsici* BCSIR-Raj-01 was comprehensively annotated using both the EDGE Bioinformatics platform and the PATRIC Comprehensive Genome Analysis pipeline. The assembly produced a high-quality draft genome with a total size of 4,620,335 bp across 37 contigs in PATRIC (Figure 1a). The G+C content was consistent across platforms, averaging 37.17%. Assembly metrics indicated high contiguity, with an N50 of 950,813 bp and an L50 of 2, reflecting robust sequencing depth and assembly fidelity. Genome annotation revealed extensive coding potential. PATRIC reported 4,699 CDSs, of which 1,443 were annotated as hypothetical (Table 1). These findings underscore a substantial fraction of the genome comprising uncharacterized or strain-specific genes, suggesting opportunities for novel functional discoveries. In PATRIC, 938 CDSs were assigned EC numbers, 801 were associated with Gene Ontology (GO) terms, and 708 were linked to KEGG pathways, revealing key roles in central metabolism, environmental adaptation, and secondary metabolite biosynthesis. Additionally, PATRIC identified 4,331 genus-specific protein families (PLFams) and 4,384 cross -genus protein families (PGFams), highlighting conserved and divergent functional traits across the *Lysinibacillus* genus. Phylogenomic reconstruction using the Type Strain Genome Server (TYGS) further clarified evolutionary relationships. BCSIR-Raj-01 was grouped in a clade with *L. capsici* and *L. boronitolerans*, supported by moderate branch lengths, indicating a recent common ancestor and high genetic similarity (Figure 1b). This clade was distinct from those containing *L. sphaericus* and other members of the genus, corroborating findings from ortholog clustering. Whole-genome similarity analysis based on average nucleotide identity (ANI) revealed that *L. capsici* BCSIR-Raj-01 shares the highest similarity with *L. capsici* strain anQ-h6 (96.90%) and *L. boronitolerans* JCM21713 (95.10%), supporting its classification within the genus. Distant similarities were observed with *L. agricola* FJAT-51161 (80.92%) and *L. alkalisoli* CGMCC1.15760 (70.62%). Surprisingly, high ANI values were also recorded with *L. sphaericus* DSM28 (99.18%) and *L. tabacifolii* KCTC33042 (99.15%), indicating close evolutionary relationships (Figure 1c).

**Table 1:** Genome assembly and annotation statistics for *Lysinibacillus capsici* BCSIR-Raj-01.

| Parameter | Value |
| --- | --- |
| Contigs | 37 |
| GC Content (%) | 37.17 |
| Genome Length (bp) | 4,620,335 |
| Contig N50 (bp) | 950,813 |
| Contig L50 | 2 |
| CDS | 4,699 |
| tRNA genes | 28 |
| rRNA genes | 3 |
| Hypothetical proteins | 1,443 |
| Proteins with functional assignments | 3,256 |
| Proteins with EC number assignments | 938 |
| Proteins with GO assignments | 801 |
| Proteins mapped to KEGG pathways | 708 |
| Proteins assigned to PATRIC PLFams | 4,331 |
| Proteins assigned to PATRIC PGFams | 4,384 |

**Figure 1.**
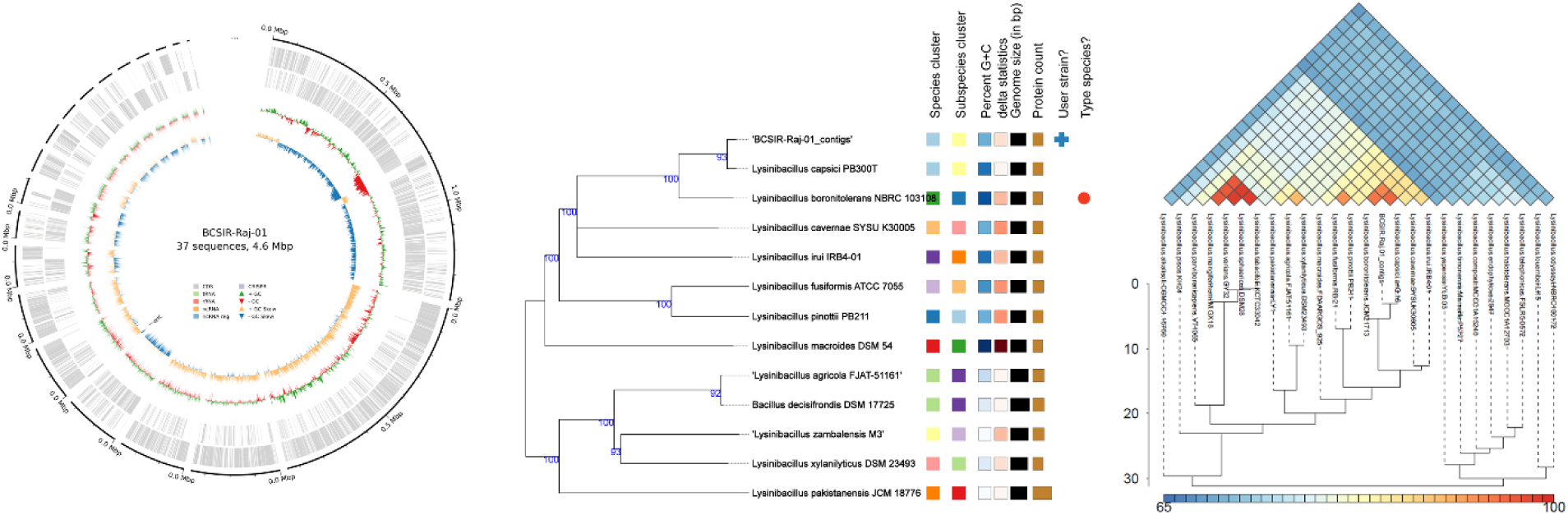
Genome assembly, functional annotation, and comparative genomic analysis of *Lysinibacillus capsici* BCSIR-Raj-01. **(a)** Circular genomic view of *L. capsici* BCSIR-Raj-01 generated by Bakta. **(b)** Phylogenetic tree of *Lysinibacillus* strains constructed using the Type Strain Genome Server (TYGS). **(c)** Average Nucleotide Identity (ANI) of *L. capsici* BCSIR-Raj-01 compared to other *Lysinibacillus* strains. The color bar on the right of the dendrogram represents ANI value. Scale under the dendrogram shows the distance which reflects the degree of similarity/difference in ANI between the strains.

#### Functional genome of *Lysinibacillus capsici* BCSIR-Raj-01

The genome of *L. capsici* BCSIR-Raj-01exemplified a diverse and wide-ranging metabolism. It contains 200 carbohydrate metabolism, 300 amino acid metabolism, and 122 nucleotide synthesis genes, which represent high carbon, nitrogen, and nucleotide turnover (Figure 2a). Genes for DNA repair (68), cofactor/vitamin biosynthesis (155), membrane transport (59), and iron acquisition (31) ensure genomic stability and flexibility under varying conditions. Stress response (39 genes) and defense systems (47 genes) were identified, but no major virulence factors were found to suggest an environmental, not pathogenic, function.

**Figure 2.**
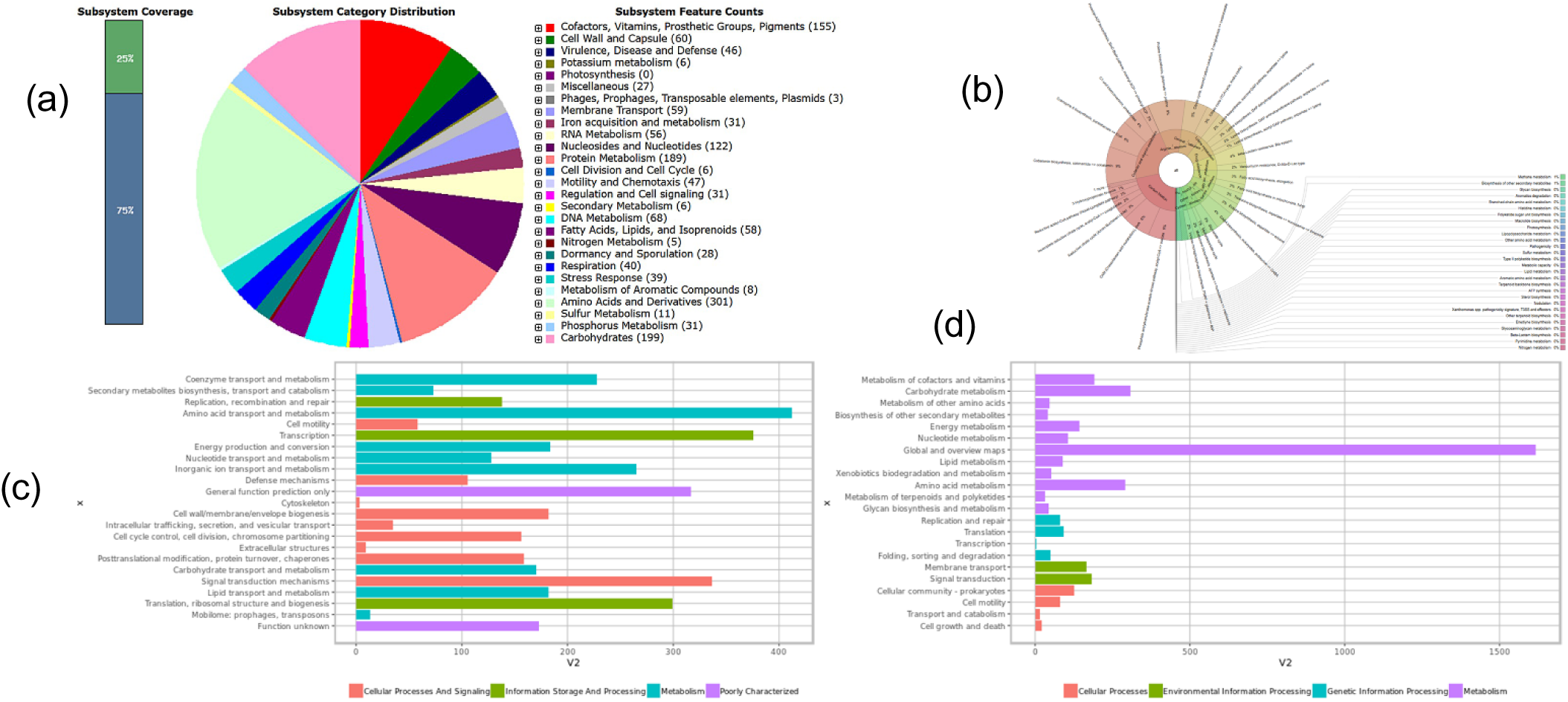
Genomic functional annotation of *Lysinibacillus capsici* BCSIR-Raj-01. **(a)** Subsystem distribution of *L. capsici* BCSIR-Raj-01 based on RAST SEED analysis. The pie chart organizes the presented subsystems by cellular process, and the number of protein-coding genes that are predicted to be involved in that cellular process are indicated. **(b)** KronaPlot of the Metagenome Assembled Genomes (MAGs) with functional annotation of *L. capsici* BCSIR-Raj-01 sample. The metabolic function is classified by systemic hierarchy levels, from higher levels in the center of the chart progressing outward until metabolic level. (c) Clusters of Orthologous Groups (COG) functional classification. (d) Kyoto Encyclopedia of Genes and Genomes (KEGG) pathway assignment.

Specialized function pathways included lipid metabolism (58 genes), signal transduction (32), sulfur (11), and phosphorus metabolism (33), whereas aromatic compound degradation only required 8 genes, indicating compromised xenobiotic processing. Key pathways such as glycolysis, the TCA cycle, oxidative phosphorylation, and amino acid biosynthesis were preserved (Figure 2b). Motility (chemotaxis, flagella) and mobile genetic elements (transposases, integrases, plasmid proteins) genes suggest potential for colonization in the environment and adaptability. COG analysis categorized genes in 22 groups (Figure 2c). The most common were Amino acid transport and metabolism (412 genes), Transcription (376), and Signal transduction (337). It also had considerable representation in Translation and ribosomal biogenesis (299), Energy production (184), and Inorganic ion transport (265). Surprisingly, 173 genes were of unknown function, indicating scope for new discoveries.

KEGG annotation showed metabolism predominance (2,917 genes), highlighting carbohydrate (309), amino acid (290), and cofactor/vitamin metabolism (190) particularly (Figure 2d). Energy metabolism (142) and xenobiotic degradation (52) genes show strong biosynthetic and catabolic potential. Environmental adaptation manifested through membrane transport (165) and signal transduction (183) genes, whereas genetic information processing pathways were translation (91) and replication/repair (80). Interestingly, antimicrobial resistance (38) and infectious disease: bacterial (27) genes are more indicative of intrinsic defense functions and less of pathogenicity. Together, these results reveal that *Lysinibacillus capsici* BCSIR-Raj-01 is a stress-tolerant and metabolically versatile bacterium with a high degree of stress response, adaptation, and nutrient cycling ability, yet with a largely non-pathogenic profile.

#### Pangenome Structure and Evolutionary Features of *Lysinibacillus capsici* BCSIR-Raj-01

To characterize the genomic novelty and evolutionary positioning of *Lysinibacillus* BCSIR-Raj-01, comparative pan-genome analysis was performed with 25 publicly available *Lysinibacillus* genomes using the iPGA platform. Unique genes identified in BCSIR-Raj-01 were categorized based on Clusters of Orthologous Groups (COG) and were linked to diverse biological processes, suggesting niche-specific adaptations. Notably, genes such as *wcaA* (involved in cell surface polysaccharide biosynthesis) and *mcrB* (DNA restriction-modification system) may contribute to structural resilience and phage defense. The presence of *pglC*, a glycosyltransferase associated with cell wall biosynthesis, and *mcrC*, involved in carbon fixation, implies potential roles in environmental stress tolerance and metabolic flexibility. Additional genes such as *btuD_8* (NADH dehydrogenase subunit), *nrdI_2* (transcriptional regulator), and *ihfA* (ribosomal RNA methylation) further support a broad metabolic and regulatory repertoire. Genes encoding *bshB2_1* (beta-lactamase) and *ykfC* indicate putative antibiotic resistance, while *xerC*, encoding a protein with a cyclic nucleotide-binding domain, suggests unique signal transduction capacities (Figure 3a).

**Figure 3.**
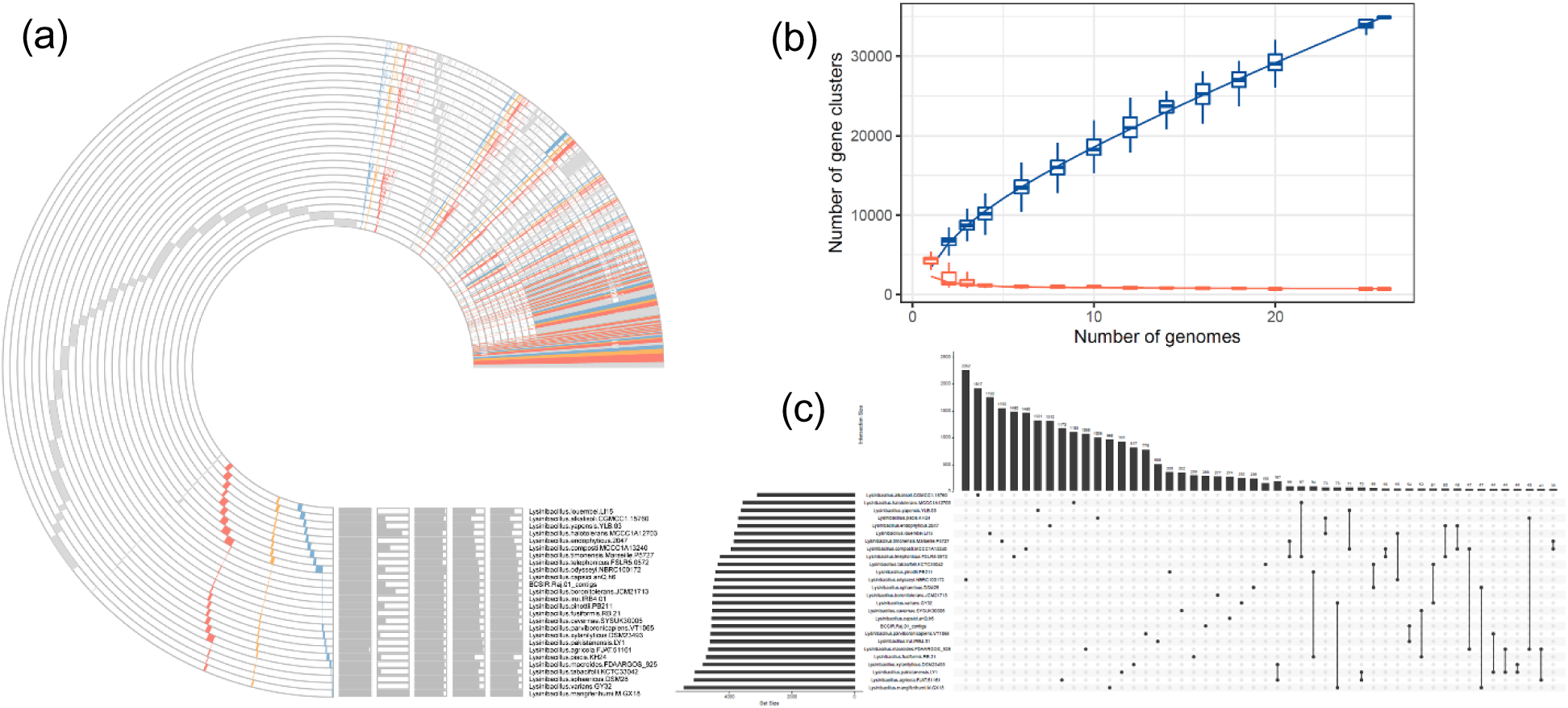
Functional categorization, phylogenetic similarity, and pangenome analysis of *Lysinibacillus capsici* BCSIR-Raj-01. **(a)** COG annotation showing the core gene clusters, unique gene clusters, number of contigs, GC content (GC), genome length and genome completeness in context of metabolism, information storage and processing, cellular processes and signaling and poorly characterized gene cluster. Different colors indicate the functional classification of the different gene clusters. The range of red arc represents the core gene clusters and the range of purple arc represents the unique gene clusters among the strains. **(b)** An UpSet plot representation of the shared and unique gene families between 25 *Lysinibacillus* species and *L. capsici* BCSIR-Raj-01. The horizontal bars (left) show the total number of gene families in each species; vertical bars represent the frequency for each intersection, and colored circles highlight the species that are part of the intersection. **(c)** Rarefaction curves of the accumulations of pan genes and core genes. The horizontal axis represents the number of samples selected; the vertical axis represents the number of genes in the selected sample combinations.

These results collectively support the taxonomic placement of BCSIR-Raj-01 within the *Lysinibacillus* genus while highlighting its distinct genomic traits. To visualize shared and strain-specific genes, an UpSet plot was generated for the analyzed *Lysinibacillus* genomes. The largest intersection cluster, containing 2,262 genes, represented the conserved core shared among the majority of strains (Figure 3b). Decreasing intersection sizes revealed progressively more strain-specific gene clusters, indicating functional divergence and the presence of species-unique adaptive traits. Notably, BCSIR-Raj-01 harbored a set of unique genes not found in other genomes, suggesting distinct ecological strategies or metabolic capabilities. Pangenome analysis revealed an open genomic architecture within the *Lysinibacillus* genus. The total pangenome expanded from 3,123 genes with a single genome to 32,637 genes upon inclusion of all 26 genomes, indicating continuous discovery of novel genes with each added strain. In contrast, the core genome decreased to just 689 genes as genome numbers increased, reflecting substantial genomic variability and species diversification (Figure 3c). The relatively small core genome, coupled with a large accessory component, supports the hypothesis of high functional plasticity and adaptive evolution within the genus.

#### Comparative metabolic profiling of Lysinibacillus BCSIR-Raj-01

Comparative analysis using OrthoVenn3 with *Lysinibacillus sphaericus* DSM 28 and *Lysinibacillus capsici* NEB659 provided insights into strain-level genomic differentiation. *Lysinibacillus capsici* BCSIR-Raj-01 possessed 15 unique gene clusters not shared with the other strains, whereas *L. sphaericus* DSM 28 and *L. capsici* NEB659 harbored 17 and 83 unique clusters, respectively (Figure 4a). These unique clusters likely encode strain-specific adaptations, potentially related to host association or ecological niche specialization. Despite the presence of unique genes, a substantial proportion of the gene content was conserved. *L. capsici* NEB659 and *L. capsici* BCSIR-Raj-01 shared 835 orthologous gene clusters, suggesting close evolutionary proximity. In contrast, *L. capsici* BCSIR-Raj-01 shared only 77 clusters exclusively with *L. sphaericus* DSM 28, indicating greater phylogenetic distance from this strain. All three genomes shared a core set of 3,056 clusters, likely representing essential housekeeping genes and conserved pathways within the genus. Comparative analysis of functional protein families between *L. capsici* BCSIR-Raj-01, *L. capsici* NEB659, and *L. sphaericus* DSM 28 revealed a conserved core of nutrient assimilation pathways, including genes encoding glutaminase, glutamine synthetase (nitrogen metabolism), sulfite reductase and sulfate adenylyltransferase (sulfur assimilation), and polyphosphate kinase and ATP synthase (phosphorus metabolism) (Figure 4b). However, inter-strain variability in enzyme isoforms and gene copy number was evident. For example, *L. capsici* BCSIR-Raj-01 uniquely encodes multiple variants of cysteine desulfurase, suggesting enhanced sulfur metabolism flexibility. Further differences were observed in fatty acid and cofactor biosynthesis, where *L. capsici* BCSIR-Raj-01 harbored strain-specific enzymes such as octanoate-acyl carrier protein transferase, potentially conferring ecological advantages under nutrient-limited conditions. Divergence in purine metabolism and folate biosynthesis genes, particularly in *L. sphaericus* DSM 28 and *L. capsici* NEB659, reflect niche-specific adaptations and variation in nucleotide salvage capabilities (Figure 4c). Although all three strains shared several conserved antibiotic resistance determinants, strain-specific resistance genes were also detected, suggesting unique strategies to withstand environmental antimicrobial pressures.

**Figure 4.**
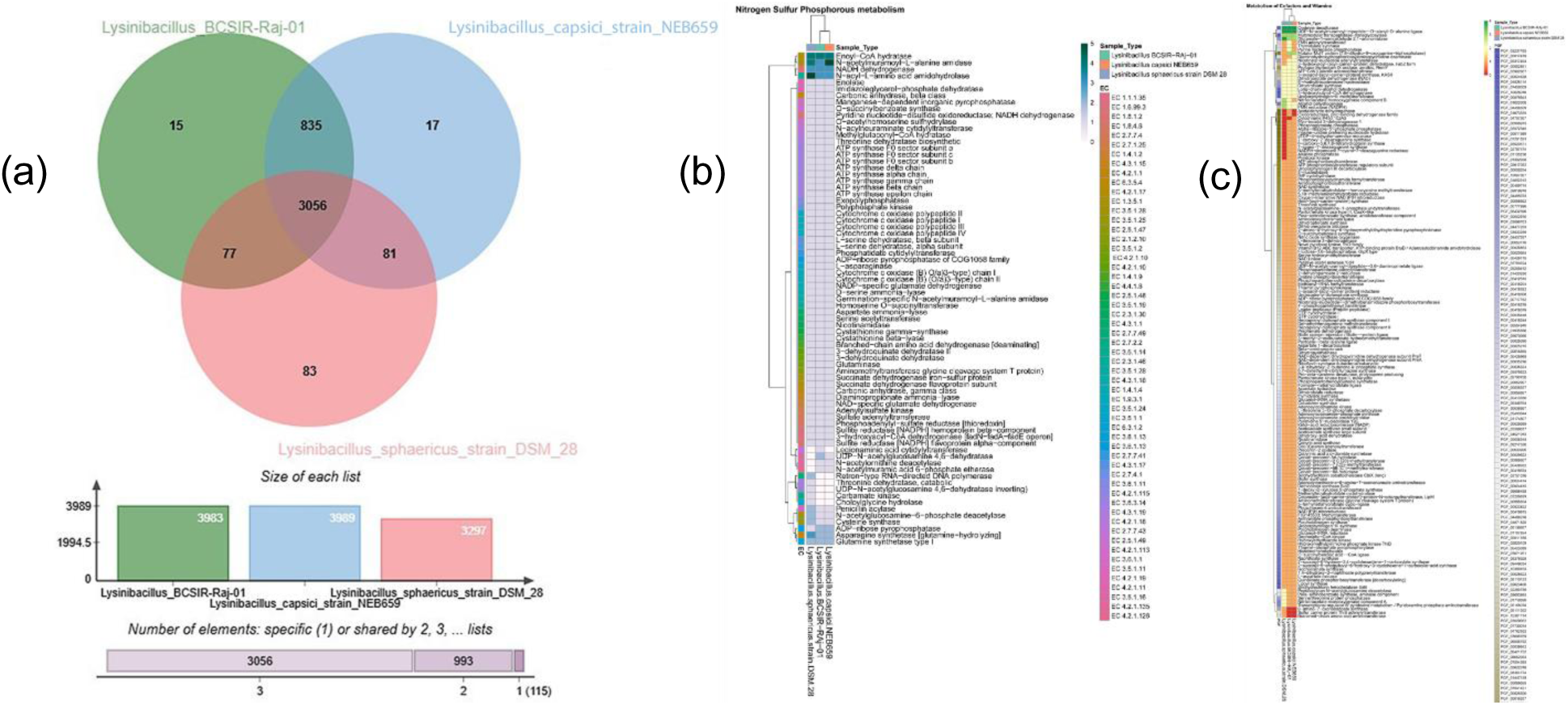
Metabolic capabilities, stress response, and comparative protein family analysis of *Lysinibacillus capsici* BCSIR-Raj-01. **(a)** Venn diagram showing the distribution of gene clusters between *L. capsici* BCSIR-Raj-01, *Lysinibacillus sphaericus* DSM 28, and *Lysinibacillus capsici* NEB659. **(b)** Comparative analysis of nitrogen, sulfur, and phosphorus metabolic pathways among *Lysinibacillus* BCSIR-Raj-01, *Lysinibacillus capsici* NEB659, and *Lysinibacillus sphaericus* DSM 28. The color of each small square represents the level of metabolite expression. **(c)** Comparative analysis of metabolism of cofactors and vitamins pathway *L. capsici* BCSIR-Raj-01, *L. capsici* NEB659, and *L. sphaericus* DSM 28. The color of each small square represents the level of metabolite expression.

#### Xenobiotic degradation, and secondary metabolism

*L. capsici* BCSIR-Raj-01 possesses a robust genetic toolkit for degrading environmental pollutants. It harbors an enriched set of alcohol and aldehyde dehydrogenases compared to its relatives, suggesting enhanced potential for short-chain hydrocarbon utilization and aldehyde detoxification (Figure 5a). Key genes for aliphatic (*lipA, adh, ald*) and aromatic hydrocarbon degradation (*catA, catE, hpaH, ndoA*) were identified, along with genes for synthetic polymer breakdown (*nylA, nylB*). Complementing this, a broad repertoire of multidrug resistance (MDR) efflux pumps (*actP, adeC, mepA, EmrAB*) and regulators (*qacE, qacR, bmrR, marR*) was present, likely conferring tolerance to antimicrobials and biocides (Figure 5b, c).

**Figure 5.**
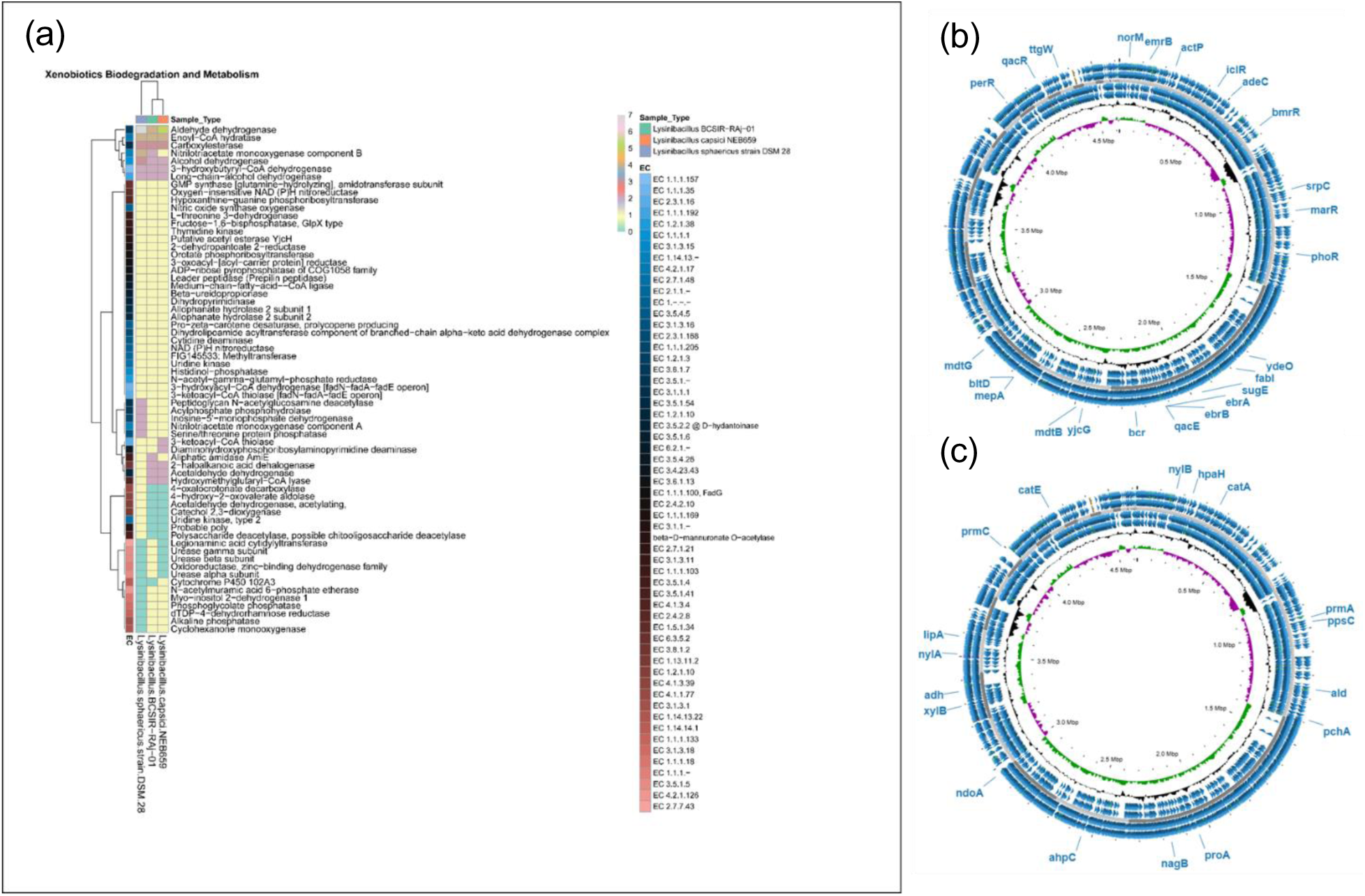
Comparative analysis of protein families involved in xenobiotic biodegradation and metabolism across *Lysinibacillus capsici* BCSIR-Raj-01. (a) Comparative analysis of xenobiotic biodegradation and metabolism pathways among *L. capsici* BCSIR-Raj-01, *Lysinibacillus capsici* NEB659, and *Lysinibacillus sphaericus* DSM 28. The color of each small square represents the level of metabolite expression. (b) Circular genome representation of BCSIR-Raj-01 showing different biocide resistant genes. In the cycle diagram, the first circle indicates the GC content, and the second circle represents the GC-Skew value. The white and gray arrows in the middle represent the open reading frames (ORFs) of the genome. The second and third circles represent CDS on the positive and negative strands, respectively. (c) Circular genome representation of *L. capsici* BCSIR-Raj-01 showing different Hydrocarbon degradation genes.

The strain’s biocontrol potential is underscored by its capacity for secondary metabolite production. Comparative analysis showed conserved enzymes for foundational biosynthesis pathways across all three strains (Figure 6a). *L. capsici* BCSIR-Raj-01 was distinguished by an elevated number of acetyl-CoA carboxyl transferase subunits and unique genes like 1-deoxy-D-xylulose 5-phosphate synthase (terpenoid biosynthesis) and dihydropteroate synthase (folate pathway). Bioinformatic mining predicted nine biosynthetic gene clusters (BGCs). BAGEL4 identified clusters for a bacteriocin (Propionicin_SM1), a lasso peptide, and a lantibiotic (LAPS) (Figure 6b). antiSMASH predicted additional clusters for terpenes, T3PKS, β-lactones, NRPS-like systems, and hybrid pathways, indicating a strong potential to produce diverse antimicrobial and bioactive compounds (Figure 6c).

**Figure 6.**
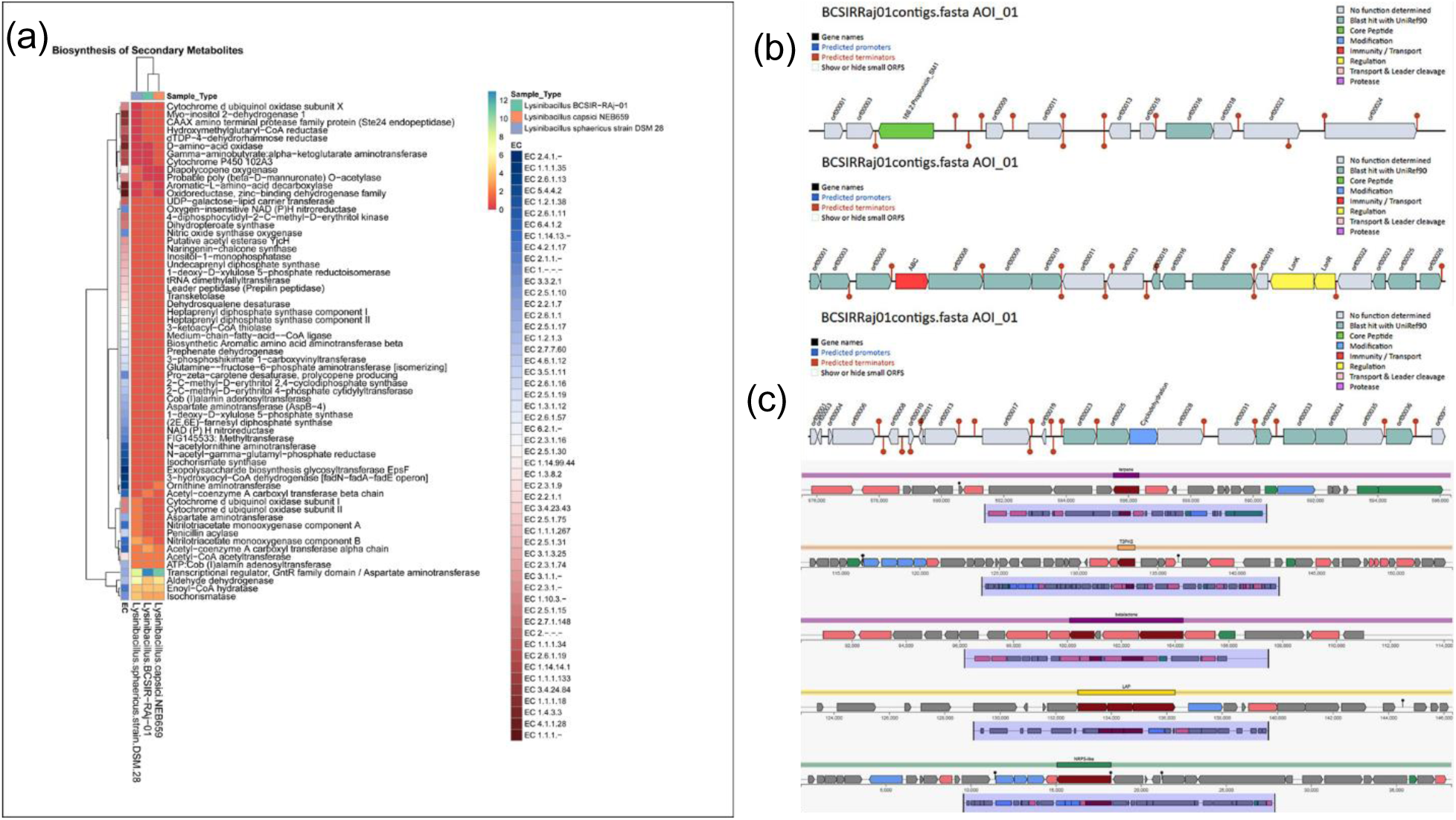
Comparative analysis of secondary metabolite biosynthesis pathways in *Lysinibacillus* strains and gene cluster identification in *Lysinibacillus capsici* BCSIR-Raj-01. (a) Comparative analysis of secondary metabolite biosynthesis pathways among *L. capsici* BCSIR-Raj-01, *Lysinibacillus capsici* NEB659, and *Lysinibacillus sphaericus* DSM 28. The color of each small square represents the level of metabolite expression. **(b)** BAGEL4 analysis identified three distinct gene clusters in *Lysinibacillus* BCSIR-Raj-01, highlighting its potential for secondary metabolite production. **(c)** antiSMASH analysis of *L. capsici* BCSIR-Raj-01 identified six distinct gene clusters, further supporting the strain’s capacity for diverse secondary metabolite biosynthesis.

#### Stress tolerance

*L. capsici* BCSIR-Raj-01 is genetically equipped to withstand various environmental stresses. All compared strains shared core stress defense systems (e.g., catalase KatE, chaperones DnaJ/K, universal stress proteins) (Supplementary 1a). BCSIR-Raj-01 showed a heightened arsenal for metal resistance, particularly arsenic (*arsA, arsD*), with additional determinants for cadmium, copper, cobalt-zinc-cadmium (*czcD*), chromate, manganese, nickel, and zinc (Supplementary 1b). It also encoded more cold shock proteins and an additional arsenate reductase. Antibiotic resistance profiling revealed a multi-mechanism resistome including *vanW/Y* (glycopeptides), *qacJ* (biocides), *msr(G)* (macrolides), and *FosBx1* (fosfomycin) (Supplementary 1c).

#### Plant growth-promoting traits

Genome annotation showed that *L. capsici* BCSIR-Raj-01 harbors the complete set of genes (highlighted in yellow) for the shikimate pathway, enabling the conversion of phosphoenolpyruvate (PEP) and erythrose-4-phosphate into chorismate. Chorismate functions as a central branching metabolite for the biosynthesis of the aromatic amino acids tryptophan, phenylalanine, and tyrosine (Figure 7a). The isolate encodes the full tryptophan biosynthetic route (chorismate → anthranilate → indole-3-glycerol phosphate → L-tryptophan), confirming its ability to synthesize tryptophan de novo. Genes involved in downstream metabolism of tryptophan to indole were also identified, suggesting a capacity for producing indole-derived secondary metabolites. These branches provide precursors for plant growth–promoting traits, including indole-3-acetic acid (IAA) biosynthesis (Figure 7b) and siderophore production (Figure 7c2), both of which were validated experimentally. In addition, genes for phenylalanine and tyrosine biosynthesis were present, indicating broader potential for aromatic secondary metabolite production. Collectively, these genomic features underpin the observed plant growth–promoting and biocontrol functions of *L. capsici* BCSIR-Raj-01.

**Figure 7.**
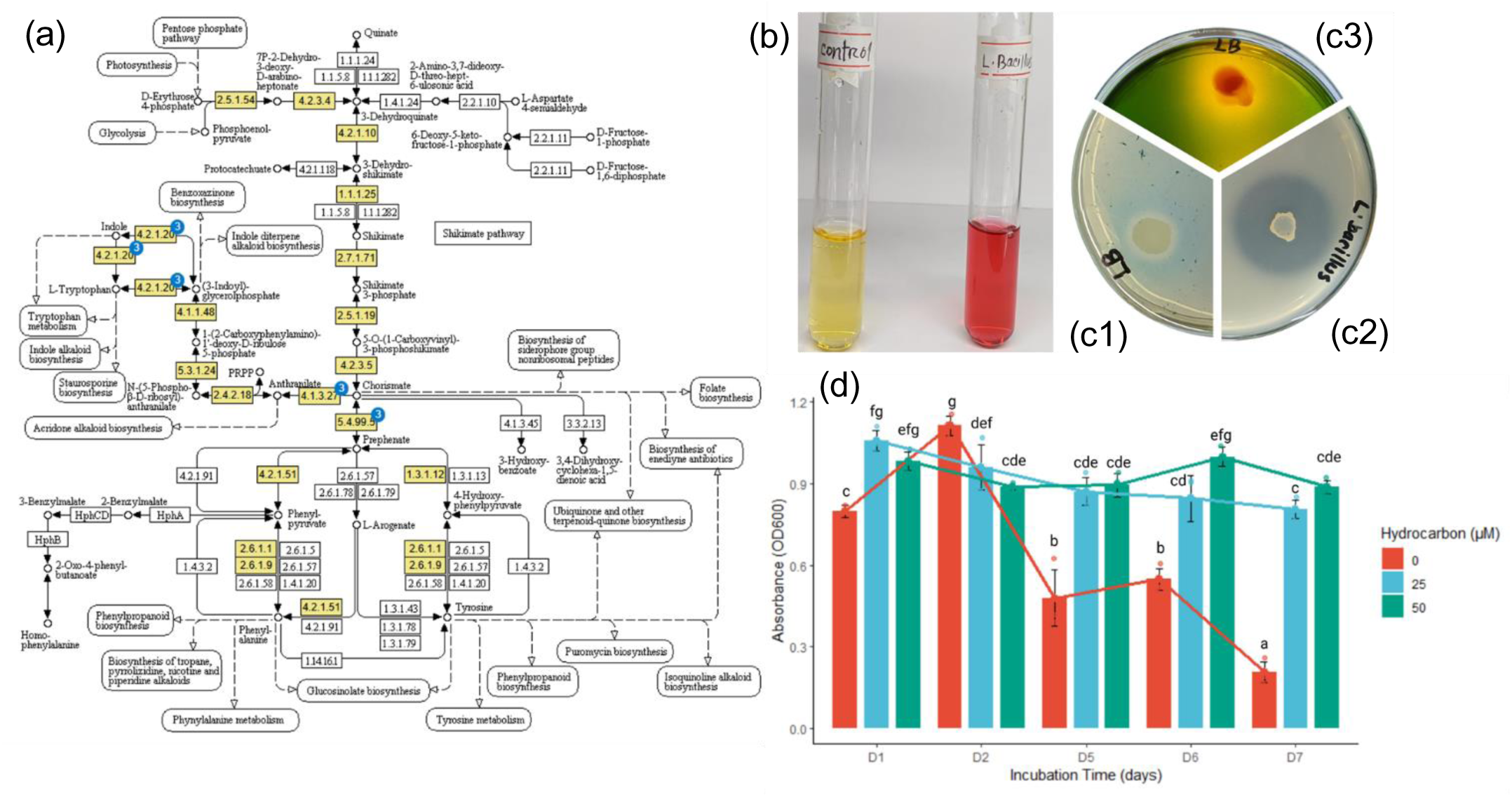
Plant growth-promoting activities and Hydrocarbon Utilization potential of *Lysinibacillus capsici* BCSIR-Raj-01. **(a)** Schematic Representation of the Tryptophan Biosynthesis Pathway in *L. capsici* BCSIR-Raj-01. This figure illustrates the complete tryptophan biosynthesis pathway identified in the genome of *L. capsici* BCSIR-Raj-01. The yellow boxes highlight the presence of genes encoding enzymes that catalyze each step of the pathway, indicating that the organism possesses the genetic potential to synthesize tryptophan. **(b)** Indole-3-acetic acid (IAA) production by *L. capsici* BCSIR-Raj-01 in an L-tryptophan amended medium was significant and comparable to the IAA levels produced by *E. coli* ATCC. IAA biosynthesis, typically occurring via the tryptophan-dependent pathway, can proceed through multiple intermediates, including tryptophol, tryptamine, indole-3-pyruvic acid, and indole-3-acetamide pathways. The exact biosynthetic route in *L. capsici BCSIR-Raj-01* remains to be elucidated. **(c1)** Phosphate solubilization activity of *L. capsici* BCSIR-Raj-01 on Pikovaskaya’s agar medium demonstrated moderate solubilization, forming a clear halo around the colony, indicative of the strain’s capacity to convert insoluble phosphate into a soluble form. **(c2)** Siderophore production by *L. capsici* BCSIR-Raj-01 was observed on CAS agar medium, which produced the highest quantity of siderophores. Detection of Nitrogen Fixation Activity in *L. capsici* BCSIR-Raj-01 Using Nitrogen-Free Medium. **(c3)** A yellow color surrounding the bacterial colonies indicates nitrogen fixation coupled with acid production, likely due to carbon source utilization. **(d)** Mean ± SE (n=4) from the two-way ANOVA of absorbance indicating *Lysinibacillus* growth among the tested hydrocarbon concentration on different incubation time. Each dot shows individual replications measured absorbance. The coloured lines connecting the top of each bar indicate the trend of growth. The different letter suggests a significantly different group (Multiple R-squared (r^2^) = 0.94, p<0.001, ANOVA, Tukey HSD)

On Pikovskaya’s agar, *L. capsici* BCSIR-Raj-01 formed clear halos around colonies, indicating the ability to solubilize inorganic phosphate (Figure 7c1).

Genomic analysis of *L. capsici* BCSIR-Raj-01 revealed multiple genes implicated in inorganic and organic phosphate metabolism. Several loci encoding alkaline phosphatases (*phoA*), phosphohydrolases, and the *phn* operon (*phnA, phnW, phnX*) were identified, supporting the strain’s capacity to utilize diverse phosphonate substrates (Figure 8). In addition, genes for polyphosphate kinase (*ppk*), exopolyphosphatase (*ppx*), and phosphate transporters (*phoU*, *phoP*, *phoR*) were present, suggesting a robust genetic basis for phosphate acquisition and homeostasis.

**Figure 8:**
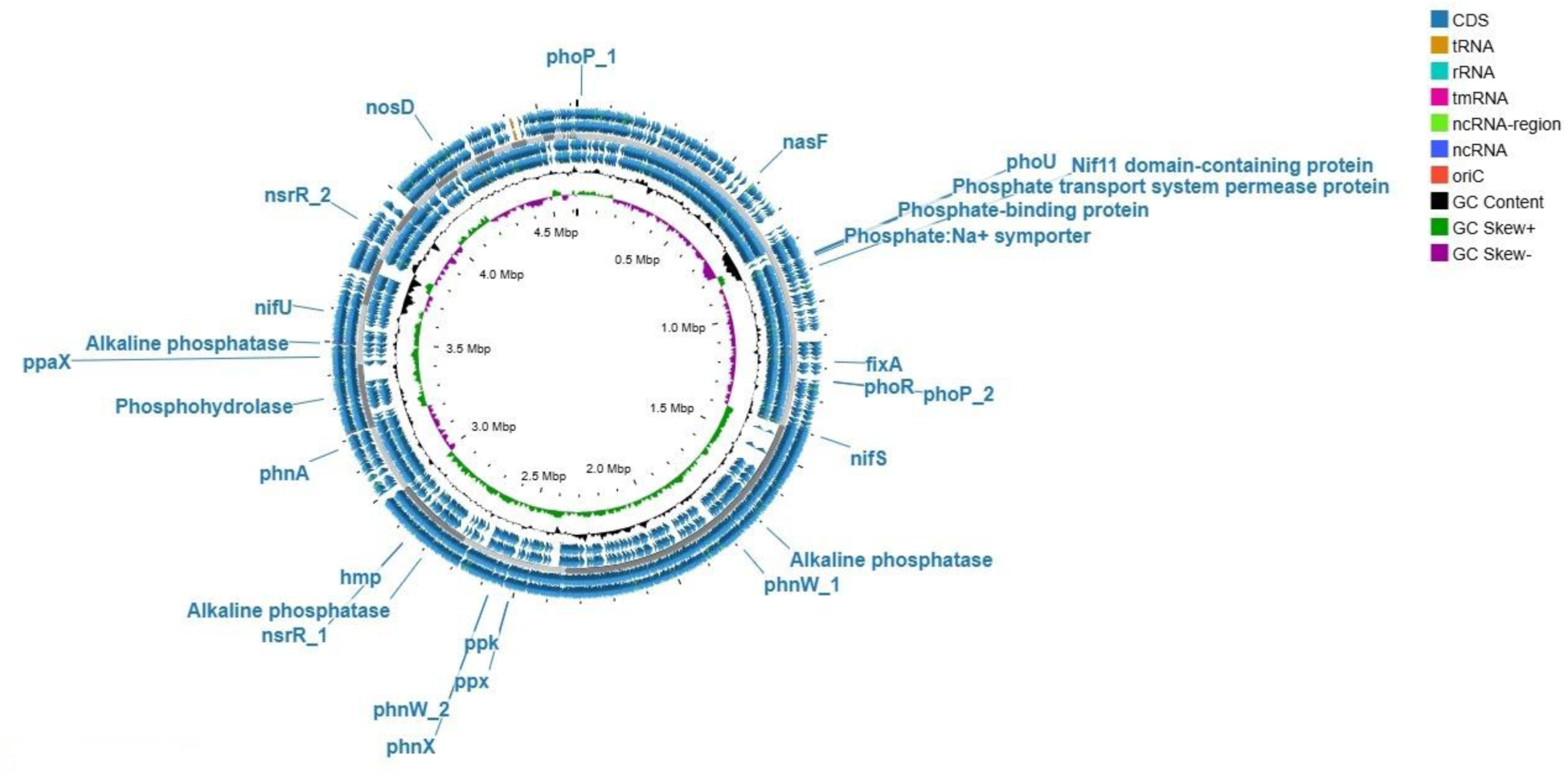
Genome organization of *Lysinibacillus capsici* BCSIR-Raj-01 highlighting nitrogen and phosphorus metabolism genes. Circular genome map of *Lysinibacillus capsici* BCSIR-Raj-01 showing genes associated with nitrogen and phosphorus metabolism. Outer rings represent coding sequences (forward and reverse strands), followed by tRNA, rRNA, and ncRNA/tmRNA regions. Inner rings depict GC content (green) and GC skew (purple/black). Key genes include *nifU*, *nifS*, *fixA*, *hmp*, *nsrR*, *nosD*, *phoP*, *phoR*, *phoU*, *ppaX*, and *phnX*, reflecting the strain’s potential for nitrogen assimilation and phosphate utilization.

*Lysinibacillus capsici* BCSIR-Raj-01 exhibited robust growth on nitrogen-free medium, accompanied by a distinct color shift of bromothymol blue from green to yellow, indicative of medium acidification and consistent with proton release during ammonium assimilation under diazotrophic metabolism (Fig. 7c3). Genome analysis identified several nitrogen metabolism–associated loci, including *nifU*, *nifS*, and *fixA*, although the canonical *nifHDK* operon was not confirmed (Figure 8). In addition, the genome harbored genes linked to nitrogen metabolism (*nsrR*, *nasF*, *hmp*) and the denitrification-associated assembly factor *nosD*, suggesting auxiliary involvement in nitrogen cycling. However, the absence of *nosZ*, the only catalytic gene for N₂O reduction, as well as *nap/nar* and *nirS/nirK* indicates that this strain lacks the capacity for canonical denitrification.

#### Phenotypic validation: hydrocarbon utilization and larvicidal activity

Phenotypic assays confirmed genomic predictions. A two-way ANOVA revealed that bacterial growth was significantly influenced by incubation time (p<0.001), hydrocarbon concentration (p<0.001), and their interaction (p<0.001) (Table 2). In a hydrocarbon-free control, growth peaked at Day 2 (OD₆₀₀=1.15) before sharply declining, indicating nutrient exhaustion. In contrast, cultures amended with 25 µM or 50 µM diesel hydrocarbon sustained stable growth over 7 days, with the 50 µM treatment showing a slight peak at Day 6 (OD₆₀₀=1.07) (Figure 8d). From Day 5 onward, both hydrocarbon-treated groups exhibited significantly higher growth than the control (p<0.001), confirming the strain’s ability to utilize hydrocarbons as a carbon source.

**Table 2:** Two-way ANOVA results for the effects of incubation time and added hydrocarbon concentration on *Lysinibacillus* growth.

| Response | Absorbance indicating *Lysinibacillus* growth |  |  |  |
| --- | --- | --- | --- | --- |
|  | Sum Sq | Df | F value | Pr(>F) |
| Incubation Time (T) | 1.01202 | 4 | 94.871 | $< 2.2\text{e-}16$ |
| Added hydrocarbon (C) | 1.12325 | 2 | 210.597 | $< 2.2\text{e-}16$ |
| T X C | 1.08263 | 8 | 50.745 | $< 2.2\text{e-}16$ |

The vegetative cells of BCSIR-Raj-01 exhibited potent, species-specific larvicidal activity against mosquito vectors (Table 3). *Culex quinquefasciatus* (LC₅₀ = 0.05 mgL⁻¹) and *Anopheles* sp. (LC₅₀ = 0.07 mgL⁻¹) were highly susceptible. In stark contrast, *Aedes aegypti* demonstrated marked tolerance, with an LC₅₀ value (1.9 mgL⁻¹) 38-fold and 27-fold higher than that for *Cx. quinquefasciatus* and *Anopheles* sp., respectively. This trend was amplified at the LC₉₀ level, underscoring a highly specific mode of action.

**Table 3:**
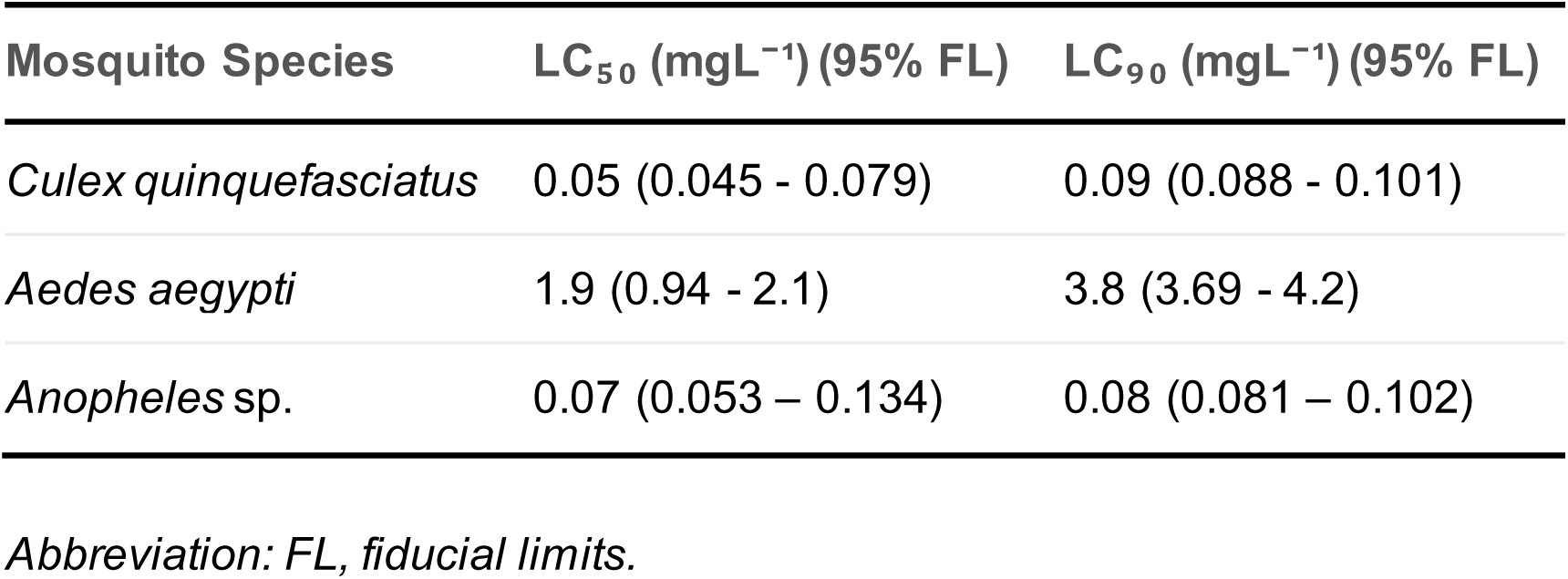
Larvicidal activity of *Lysinibacillus capsici* BCSIR-Raj-01 vegetative cells against third-instar mosquito larvae.

| Mosquito Species | LC<sub>50</sub> (mgL<sup>-1</sup>) (95% FL) | LC<sub>90</sub> (mgL<sup>-1</sup>) (95% FL) |
| --- | --- | --- |
| *Culex quinquefasciatus* | 0.05 (0.045 - 0.079) | 0.09 (0.088 - 0.101) |
| *Aedes aegypti* | 1.9 (0.94 - 2.1) | 3.8 (3.69 - 4.2) |
| *Anopheles* sp. | 0.07 (0.053 – 0.134) | 0.08 (0.081 – 0.102) |

Abbreviation: FL, fiducial limits.

#### Defense systems

The genome of BCSIR-Raj-01 is equipped for genomic plasticity and defense. We identified 72 mobile genetic elements (MGEs), predominantly involved in replication/recombination/repair (n=37) and phage-related functions (n=14) (Supplementary 2a). Two prophage regions were detected: one incomplete (24.6 kb) and one intact (52.9 kb), the latter showing similarity to PHAGE_Geobac_E3 (Supplement Figure 1b). A sophisticated defense arsenal was uncovered, including Type II and IV restriction-modification systems, an atypical CRISPR-Cas system (Cas proteins on contigs 11, 16), and additional systems like Dodola and AbiL, indicating a multi-layered strategy to resist phage predation and foreign DNA (Supplementary 2c, d).

## DISCUSSION

The multifunctionality of microbial inoculants is increasingly viewed as a cornerstone of sustainable agriculture and environmental biotechnology. Just as the introduction of synthetic fertilizers and pesticides once revolutionized food production, the next frontier depends on harnessing microbes that can simultaneously enhance plant nutrition, suppress pathogens, and mitigate pollution. In this context, our genomic and phenotypic characterization of *Lysinibacillus capsici* BCSIR-Raj-01 provides a timely example of how a single strain can integrate these functions.

Comparative genomics revealed that *L. capsici* BCSIR-Raj-01 shares the hallmark ecological versatility of the genus *Lysinibacillus* yet also harbors strain-specific genes linked to phosphate metabolism, nitrogen cycling, hydrocarbon degradation, and stress tolerance. These genomic features were not merely predictive; they aligned with phenotypic assays showing phosphate solubilization, IAA production, diazotrophic growth, diesel utilization, and potent larvicidal activity. The coherence between genotype and phenotype underscores the adaptive strategies that enable this endophyte to thrive in plant-associated niches while contributing to broader ecological processes.

Importantly, *L. capsici* BCSIR-Raj-01 illustrates both the promise and complexity of multifunctional bacteria: while its genomic features suggest the capacity to contribute nitrogen to host plants, the presence of *hmp* (flavohemoglobin) points to a role in nitric oxide detoxification that can yield N₂O under oxygen-limited conditions, underscoring the ecological trade-offs that shape microbial functions in agroecosystems. Similarly, it’s larvicidal activity—while valuable for vector control—likely relies on unconventional metabolites distinct from the classical Bin/Mtx toxins, suggesting untapped biochemical pathways. By situating *L. capsici* BCSIR-Raj-01 within these broader ecological and biotechnological contexts, our study not only expands the functional landscape of *Lysinibacillus* but also illustrates the opportunities and challenges of translating genomic potential into field applications.

### Genomic Features and Metabolic Versatility of *L. capsici* BCSIR-Raj-01

The genome of *Lysinibacillus capsici* BCSIR-Raj-01 highlights a metabolically versatile and environmentally adaptable bacterium with strong potential for biotechnology applications. Its high-quality assembly supports comparative and functional genomic analyses (50). Phylogenomics and ANI analyses place it closer to *L. capsici* than *L. sphaericus*, consistent with species-level diversification within the genus (51). Pan-genome analysis confirmed an open pan-genome, suggesting ongoing horizontal gene acquisition. Strain-specific genes encode functions related to metabolism, heavy metal resistance, xenobiotic degradation, and antibiotic tolerance, highlighting potential for survival in contaminated environments. Subsystem profiling revealed broad metabolic capacity and stress-response mechanisms, while the absence of canonical virulence genes supports a non-pathogenic, environmental lifestyle (52). Distinct sulfur and lipid metabolism pathways, unique to *L. capsici* BCSIR-Raj-01, further suggest niche specialization.

### Plant Growth–Promoting Traits

Genome mining of *L. capsici* BCSIR-Raj-01 revealed a complete tryptophan biosynthesis pathway (chorismate to tryptophan) (Figure 7a) and genes for downstream transformations to indole-3-glycerol phosphate and indole, indicating a functional tryptophan to indole metabolic branch (53). Indole is a central precursor for numerous indole-derived secondary metabolites and signaling molecules implicated in plant–microbe interactions (54). Consistent with these predictions, the isolated strain produced siderophores on chrome azurol S agar, suggesting capability to chelate ferric iron under Fe-limiting conditions, a trait that promotes rhizosphere colonization and can indirectly suppress pathogens by limiting iron availability (55). The genome of *L. capsici* BCSIR-Raj-01 encodes multiple tryptophan-dependent indole-3-acetic acid (IAA) biosynthetic routes, including the indole-3-pyruvic acid (IPyA), indole-3-acetamide (IAM), tryptamine (TAM), and tryptophol pathways. Comparative genomic studies have shown that many bacteria, including plant-associated taxa, harbor more than one tryptophan-dependent IAA pathway (56, 57). Although the precise route used by *L. capsici* BCSIR-Raj-01 remains unresolved, the IPyA pathway is likely predominant, as this route has been experimentally demonstrated as the main contributor to IAA production and plant growth promotion in *Bacillus* sp. (58, 59).

Genome annotation of *L. capsici* BCSIR-Raj-01 revealed several gene clusters and loci associated with phosphate solubilization, a key plant growth–promoting trait. Genes encoding alkaline phosphatases (*phoA, phoD, phoX*) were identified, which mediate the hydrolysis of organic phosphorus compounds. Regulatory components of the Pho regulon (*pho*B, *pho*R, *pst*SCAB transport system, and *pho*U) were also present, indicating a tightly controlled phosphate starvation response (60). Notably, unlike many well-characterized phosphate-solubilizing bacteria (PSB), *L. capsici* BCSIR-Raj-01 lacks the *pqqABCDE* operon required for pyrroloquinoline quinone (PQQ) biosynthesis. PQQ is the essential cofactor of glucose dehydrogenase (Gdh) (61), which catalyzes the oxidation of glucose to gluconic acid, a major organic acid involved in phosphate solubilization (62). The absence of this pathway suggests that the isolate relies primarily on phosphatase-mediated hydrolysis rather than organic acid secretion to mobilize phosphorus. Such a mechanism aligns with observations in other bacteria where incomplete *pqq* genes correspond to lower acid secretion and reliance on enzymatic hydrolysis of organic phosphorus (61, 63, 64).

The diazotrophic nature of *L. capsici* BCSIR-Raj-01, isolated from citrus leaves, demonstrates the potential of endophytes to contribute to host nitrogen nutrition and reduce dependence on synthetic fertilizers. This was phenotypically confirmed by robust growth on a nitrogen-free medium and acidification of bromothymol blue, a classical indicator of biological nitrogen fixation (BNF). Although consistent with a diazotrophic phenotype, genomic analysis revealed the absence of the canonical *nifHDK* operon, unlike endophytic diazotrophs reported from apple plants (65). This genotype–phenotype discrepancy highlights the complexity of diazotrophy, which is not invariably e ncoded by a contiguous *nif* operon. Prior research indicates that nitrogenase genes are taxonomically widespread, often subject to genomic rearrangements, and can be highly divergent, suggesting the potential for cryptic or alternative nitrogenase systems that evade detection by standard annotation pipelines (66–70). Furthermore, the potential involvement of plasmid-borne *nif* clusters or unassembled genomic islands remains a plausible explanation for this disparity. Therefore, definitive confirmation of N₂ fixation in *L. capsici* BCSIR-Raj-01 requires direct physiological evidence, such as ¹⁵N₂ incorporation or acetylene reduction assays, to exclude growth supported by trace environmental nitrogen. Notwithstanding the need for this confirmation, our finding of a diazotrophic *Lysinibacillus* endophyte broadens the ecological scope of this genus, which has long been recognized mainly as a rhizosphere -associated plant growth-promoting bacterium (71).

Growing evidence supports this endophytic niche; recent studies have reported endophytic *Lysinibacillus* sp., including *L. capsici* and related strains with putative nitrogen-fixing ability, from ginger (12) and maize (72). This is consistent with the earlier description of a diazotrophic, endophytic *Lysinibacillus sphaericus* from rice (73), collectively underscoring an emerging role for this genus as an endophytic diazotroph. Moreover, research in other endophytic systems has demonstrated that in planta ammonium release (74) and synergistic microbial interactions (75) can significantly enhance nitrogen transfer to the host. This suggests that the endophytic lifestyle of *L. capsici* BCSIR-Raj-01 likely facilitates more efficient nutrient delivery compared to free-living diazotrophs.

Intriguingly, the genome of *Lysinibacillus capsici* BCSIR-Raj-01 harbors a truncated nos operon, retaining *nosD* (a putative maturation/membrane component) but lacking *nosZ* (terminal reductase *nosZ* responsible for N₂O reduction to N₂) and the core reductase genes (*nap*/*nar*, *nirS*/*nirK*), indicating an inability to perform canonical denitrification (76, 77). Instead, the presence of a *hmp* homologue (encoding flavohemoglobin) suggests a role in nitric oxide detoxification (78, 79). Under microoxic or hypoxic conditions, Hmp-mediated reactions can lead to the incidental formation of N₂O as a by-product (80–82). In complex soil consortia, such organisms may indirectly influence nitrogen cycling by modulating NO levels generated by other microbes (83, 84). Thus, *L. capsici* BCSIR-Raj-01 is more likely an accessory player in nitrogen transformations rather than a direct N₂O producer.

Further genomic analysis revealed the presence of two *nsrR* regulators (*nsrR₁* and *nsrR₂*) and the transcriptional antiterminator *nasF*. NsrR proteins are NO-sensitive repressors that govern the nitrosative stress response, and their duplication may enable refined sensing of fluctuating NO levels (85, 86). Concurrently, NasF is a key regulator that derepresses assimilatory nitrate reduction operons under nitrogen-limiting conditions (87). The co-occurrence of dual *nsrR* genes with *nasF* implies an integrated regulatory network that potentially coordinates nitrate assimilation with nitrosative stress tolerance, thereby influencing the interplay between nitrogen scavenging and incomplete denitrification. Future work involving RNA-seq under nitrate limitation and nitrosative stress would be invaluable to experimentally define the roles of these regulators.

Collectively, our genomic and phenotypic analyses position *L. capsici* BCSIR-Raj-01 as a multifunctional plant growth-promoting bacterium with a repertoire of traits including IAA production, siderophore synthesis, organic phosphate solubilization, and diazotrophy. The genomic co-occurrence of diazotrophy-associated loci (e.g., *nifU*, *nifS*, *fixA*) with a truncated denitrification gene cluster (*nosD* but lacking *nosZ*) is particularly noteworthy. This genetic configuration suggests a dual ecological role: the potential to supply fixed nitrogen to plants while also contributing to nitrous oxide (N₂O) emissions as a partial denitrifier under anoxic conditions. This functional combination underscores the ecological versatility of this strain. Rather than coupling N₂O reduction to nitrogen fixation, a capacity precluded by the lack of *nosZ*, the isolate appears capable of influencing multiple steps of the nitrogen cycle simultaneously. Such multifunctionality, balancing plant nutrition with a role in greenhouse gas flux, has been documented in other rhizosphere bacteria (84, 88–91). Our findings expand the functional and ecological scope of the genus *Lysinibacillus*, traditionally regarded as a rhizosphere-associated PGPB, and highlight its potential, albeit complex, role in agricultural ecosystems.

### Hydrocarbon degradation and bioremediation

*L. capsici* BCSIR-Raj-01 maintained robust growth in hydrocarbon-amended media, consistent with its genomic repertoire for alcohol, aldehyde, naphthalene, and nylon degradation. Enrichment of aldehyde dehydrogenases suggests enhanced detoxification capacity (92), while genes encoding alcohol dehydrogenases and fatty-acid-CoA ligases point to the ability to metabolize diverse hydrocarbon-derived substrates. This metabolic versatility exceeds that of *L. capsici* and underscores its promise for petroleum and plastic remediation.

Unlike *L. sphaericus*, BCSIR-Raj-01 lacks *amoA*, indicating that hydrocarbon metabolism occurs via alternative routes rather than AMO-mediated cometabolism, which is often inefficient and inhibitory (93). Its endophytic origin suggests dual functionality, plant tolerance enhancement and in situ hydrocarbon degradation, making it a strong candidate for rhizosphere -targeted bioremediation in polluted agroecosystems (94).

### Larvicidal activity

*L. capsici* BCSIR-Raj-01 also demonstrated potent larvicidal activity, with high efficacy against *Culex quinquefasciatus* and *Anopheles* sp., but reduced activity against *Aedes aegypti*. This mirrors *L. sphaericus* activity profiles, though *L. capsici* BCSIR-Raj-01 lacks Bin and Mtx toxin genes, implying a distinct mechanism. Genomic analysis identified biosynthetic gene clusters for terpenes, type III polyketides, bacteriocins, and lantibiotics, which may collectively underpin larvicidal activity. Terpenoids and polyketides are particularly noteworthy for their insecticidal properties and potential synergism (95).

Reduced susceptibility of *Aedes* larvae may be explained by its robust detoxification enzymes and microbiome-mediated defenses (96). Nevertheless, the high potency against *Culex* and *Anopheles* highlights *L. capsici* BCSIR-Raj-01 as a promising candidate for biological vector control, particularly as resistance to conventional insecticides continues to rise (97).

### Concluding Remarks

Overall, *L. capsici* BCSIR-Raj-01 represents a multifaceted bacterium with ecological and biotechnological relevance. It combines PGP traits, hydrocarbon-degradation capacity, and larvicidal activity, supported by both genomic predictions and laboratory validation. Its non-pathogenic profile further strengthens its potential for safe application in agriculture, environmental remediation, and vector control. Future work should focus on field validation and metabolite characterization to identify the compounds responsible for larvicidal activity and hydrocarbon degradation.

## Contributions

R.H.P. and M.Z.R. contributed equally as first author.

Conceived and designed the study: M.M.H.S., R.H.P., M.Z.R; Performed the experiments and data acquisition: M.M.H.S., M.K.H., E.A., M.A.A.O; Genome sequencing : M.M.H.S., S.R.N., S.F.C.; Data analysis: S.R.N., T.C.; Sample collection and culture characterization: M.I., M.K.H., E.A., M.A.A.O; Mosquito rearing and larvicidal activities experiment: M.Z.R, N.S., N.N.H.; Provided valuable comments on the manuscript: M.A.H., S.A., T.A.B.,B.G.; Manuscript evaluation and revision: M.M.H.S., R.H.P., M.A.H., S.A., T.A.B., B.G, M.I.; All authors have read and approved the final version of the manuscript.

